# Parallel on-chip micropipettes enabling quantitative multiplexed characterization of vesicle mechanics and cell aggregates rheology

**DOI:** 10.1101/2023.10.19.562871

**Authors:** Sylvain Landiech, Marianne Elias, Pierre Lapèze, Hajar Ajiyel, Marine Plancke, Adrian Laborde, Fabien Mesnilgrente, David Bourrier, Debora Berti, Costanza Montis, Laurent Mazenq, Jérémy Baldo, Clément Roux, Morgan Delarue, Pierre Joseph

## Abstract

Micropipette aspiration (MPA) is one of the gold standards to quantify biological samples’ mechanical properties, which are crucial from the cell membrane scale to the multicellular tissue. However, relying on the manipulation of individual home-made glass pipettes, MPA suffers from low throughput and difficult automation. Here, we introduce the sliding insert micropipette aspiration (SIMPA) method, that permits parallelization and automation, thanks to the insertion of tubular pipettes, obtained by photolithography, within microfluidic channels. We show its application both at the lipid bilayer level, by probing vesicles to measure membrane bending and stretching moduli, and at the tissue level by quantifying the viscoelasticity of 3D cell aggregates. This approach opens the way to high-throughput, quantitative mechanical testing of many types of biological samples, from vesicles and individual cells to cell aggregates and explants, under dynamic physico-chemical stimuli.

## Introduction

Mechanics is ubiquitously at play in biology, from the level of cell membranes to the tissue scale. At the cell scale, response to stimuli is related to its cytoskeleton and nucleus but also strongly depends upon the deformability of its membrane^1^. Indeed, cells divide and interact with their surroundings by remodeling their cytoplasmic membranes^2^; processes of endocytosis/exocytosis involve membrane bending^3^; permeation of drugs or nanoparticles relates to the ability of lipids constituting the membrane to accommodate changes in shape. At the multicellular scale, the capacity of cell assemblies to deform and flow is a determining factor in tissue homeostasis and evolution. This idea applies for developmental biology, since embryo morphogenesis is strongly intertwined with spatiotemporal changes and heterogeneity in fluidity^4,5^. It is also an essential ingredient for pathological situations such as solid cancers: the ability of cells to deform and spread, or jam, is key in disease progression^6^. Tissue rheology can thus be envisioned as a diagnostics tool^7^, or even to assist the prognosis of metastasis^8^.

Thus, strong efforts have been made in the last decades to engineer quantitative tools assessing mechanical properties of cell membranes^9^, cells^10^, and cell aggregates^11^, often relying on analogies with soft matter as proposed in Steinberg’s pioneering work^12^, and on concepts of rheology^13^. Stress is applied either very locally by AFM probing^14^ or on the whole tissue (through magnetic nanoparticles^15^ or by parallel plate compression), to cite only a few methods. One popular technique is micropipette aspiration MPA^16–18^, both at cell and tissue scale. Measuring to what extent a vesicle, a cell, or a tissue enters a glass tube upon aspiration permits to determine mechanical properties: bending and stretching rigidity for lipid vesicles mimicking cell membranes; Young’s modulus and effective viscosity for single cells^10^; surface tension, elasticity and viscosity for 3D cell aggregates^19^. MPA is one of the gold standards because in addition to the simplicity of its principle, it is quantitative, and it probes locally a zone that can be chosen. It also enables to some extent the change of solution surrounding the sample, and it can with reasonable experimental effort be coupled to other techniques like optical tweezers. However, it requires a complex dedicated setup: microscope, micromanipulator, precise control of the pressure in the aspiration tube. The control of the physico-chemical environment in real time is tedious: it requires several micromanipulators, and the concentration of chemicals injected around the sample is non-homogeneous. Most importantly, MPA suffers from very low throughput since objects are intrinsically probed one by one, which can be limiting due to the high sample-to-sample variability that is often typical of biological systems.

Consequently, approaches to integrate micropipettes in microfluidic devices have been proposed in the very last years. They target the above-mentioned limitations by designing channels enabling parallel trapping and fluid control at the cell (or cell aggregate) scale. A design relying on 3-level fabrication has been developed ten years ago by Lee *et al*. for cells^20^, which we improved in terms of alignment for the study of Giant Unilamellar Vesicles (GUV)^21^. Boot *et al*. have recently adapted it to 3D cell aggregates^22^. While for this design microfluidics permits automation of objects injection, it is still limited in terms of throughput. More importantly, the rectangular geometry has intrinsic limitations: a quantitative analysis is complicated and some flow remains at the corner of the traps constituting the pipette, even though recent work described the different regimes of clogging rectangles with soft objects^23^. A 2-level design was used to probe the viscoelasticity of cell nuclei in parallel thanks to constrictions^24^, simpler to implement than the previous one but still not fully quantitative. In order to relate the microscopic configuration to mechanical properties, 2D geometries permitting optical access combined to rheological measurements were used to characterize cell aggregates rearrangements^25^ or vesicles prototissues^26^, but their extension to more realistic 3D tissues is far from obvious. Indeed, standard microfabrication techniques are planar, which limits the possibility to properly integrate circular traps. 3D printing technologies that are emerging are associated to prohibitive writing time for the resolution and surface quality required here. As a way to eliminate the need for fluid confinement by surfaces, virtual walls microfluidics has recently been demonstrated to characterize both cell and spheroid mechanics^27^, with quite a high throughput but limited to a global probing of objects, with a homogeneous stress.

Thus, a micropipette aspiration method, quantitative but with a higher throughput than classical MPA, is still to be developed. We describe in this paper the SIMPA technology (sliding insert micro pipette aspiration) addressing the above-mentioned requirements, both at the scales of vesicles and multicellular aggregates. It relies on the “sliding walls” proposed by Venzac *et al*.^28^, inserting sliding elements within PDMS chips. Here, rather than reconfigurability, which was the strong point raised in^28^, and which for instance permitted studying confined tissue growth^29^, we specifically exploited the particular microfabrication features of the approach. Pipettes are designed and patterned by photolithography perpendicularly to the fabrication plane of the channel in which they are inserted (see Figure 1). This way, the objects injected in a microchannel can be blocked by pipettes of chosen shape: a circular cross-section permits quantitative measurements analyzed with classical models, since deformations occur like in standard MPA. Thanks to the integration, pipettes can be operated in parallel, which increases the throughput: we demonstrate it for 7 GUVs, and for up to 23 spheroids.

**Figure 1.**
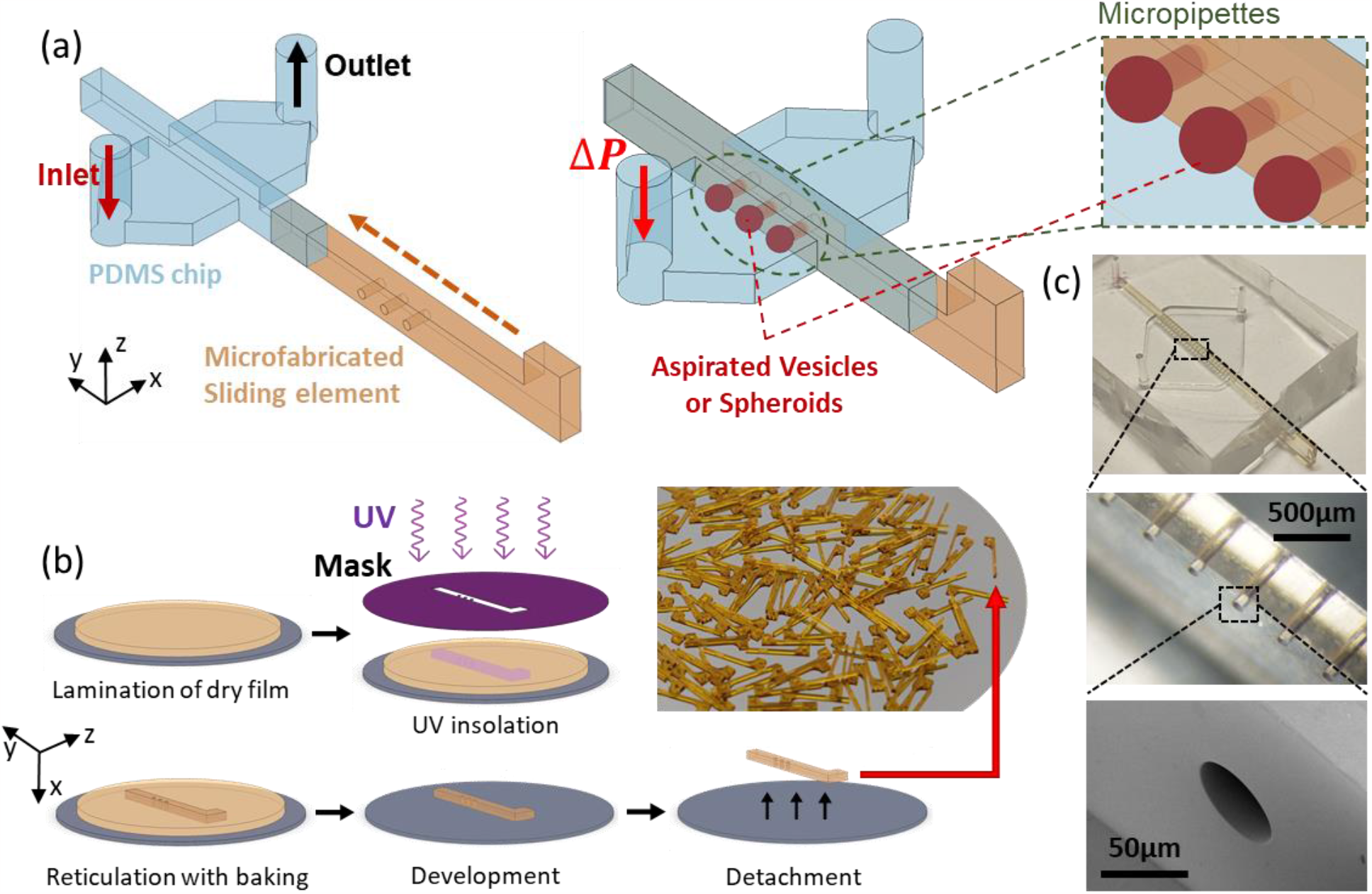
Principle: On-chip pipettes integrated in a microfluidic chip thanks to sliding elements -. (a) Parts view: PDMS chip and sliding element. Assembled view after insertion, schematic close-up of aspirated micro-objects (Giant Unilamellar Vesicles or Spheroids). b) Microfabrication workflow of the sliding elements and photograph of dozens of them, manufactured in a single batch. c) Micrographs and SEM close-ups of the pipettes integrated in the sliding elements.

In the following, we explain the design principles and fabrication technique of the chips, as well as their fluidic operation. We then demonstrate the interest of the technology by assessing two situations relevant for biophysics. First, we present the results obtained on vesicles: characterization of the elastic moduli of lipid bilayers with simple composition, study of the influence of sugar and of cholesterol on these moduli. Second, we detail the use of the devices for 3D cellular aggregates: measurements of the surface tension and viscoelastic characteristics, and study of the influence of drugs targeting cell-cell adhesion.

### Principle of the SIMPA chips: design, fabrication and operation

The microfluidic chips consisted of two parts, see Figure 1a. The fabrication protocol is detailed in the Supplementary Information. Here, we explain the main ingredients of its design, fabrication and operation.

The first part is a PDMS chip, obtained by standard soft lithography. PDMS was casted and cured on a two-level mold patterned in a photosensitive dry film (SUEX), preferred to liquid photoresist since thickness is as high as ∼500 µm. After unmolding, holes were punched in PDMS for fluidic access.

The first level of the mold corresponds to the main fluidic channel, see Figure 1(a). Its height is slightly superior to the maximum diameter of the objects to be probed with pipettes, typically ∼100 µm for studies on vesicles, and ∼450 µm for spheroids.

The fluidic configuration is quite simple with 1 input and 1 output, the channel just getting wider at the location of sliding element insertion, in order to permit objects to be trapped in parallel. Note however that the design can be complexified for additional functions: we demonstrate for instance injection of a chemical stimulus around spheroids, thanks to extra lateral channels (see Figure SI-8).

The fluidic channel is intersected by another guide, integrated in the second layer of the mold, that is open to the outside. Its purpose is to accommodate the sliding element integrating the pipettes, second part that composes our chips.

The PDMS part was then bonded by plasma on a thin layer of PDMS (50 µm), compromise between optical access for microscopy and deformation to avoid leakage.

The second element is the sliding element containing the pipettes to be integrated in the fluidic channel by insertion in the PDMS chip. This long parallelepiped including holes that constitute the pipettes was manufactured by photolithography using the same type of dry film, see Figure 1(b). After optimizing fabrication parameters, we obtained pipettes with aspect ratio up to 20 (25 µm diameter for 500 µm length) and with a low roughness: typically, only a few ∼20 nm-high asperities can be seen inside pipette, as shown in Figure 1(c) and Figure SI-1. For pipettes with lower diameters (down to 12 µm for GUVs, see Figure 2(a), and 5 µm for single cells), a multi-layer lamination protocol was used. Lamination was realized on a sacrificial layer (copper-titanium alloy), chemically etched after fabrication to release the sliding elements from the wafer. Since fabrication was realized by batches on a 4-inch wafer, up to 150 reusable SMPs (sliding micro pipettes) could be obtained in a single fabrication run, which typically took a few hours.

**Figure 2.**
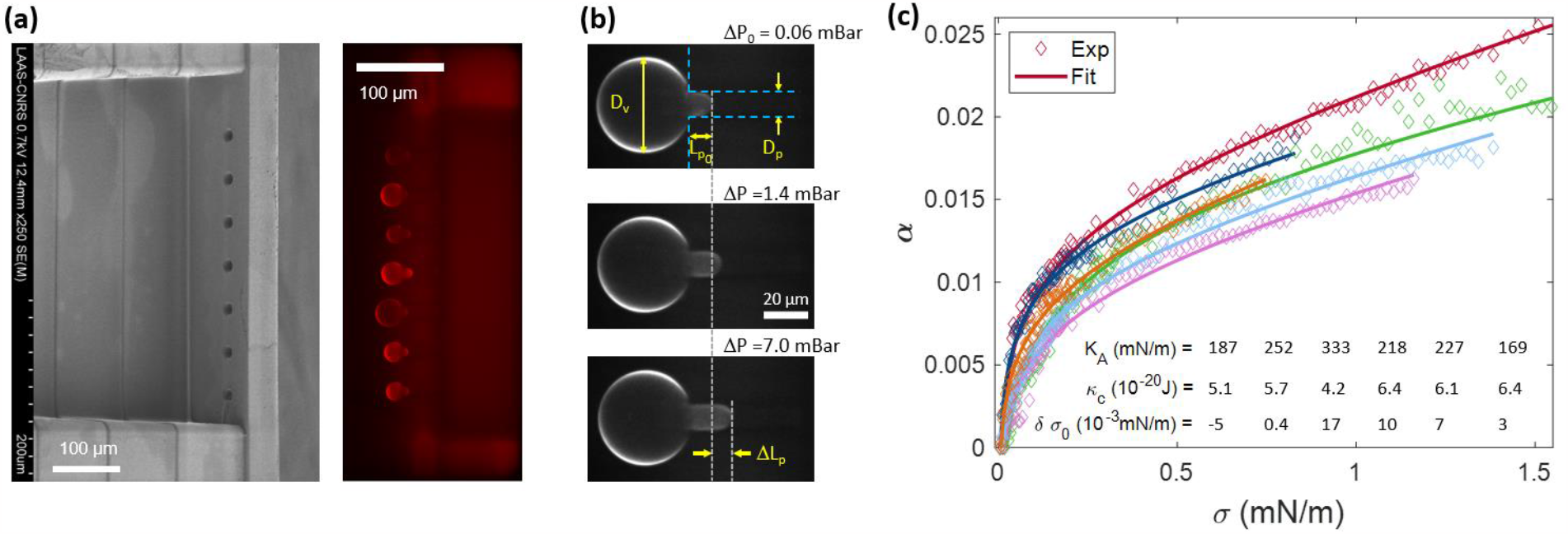
On-chip pipettes applied to Giant Unilamellar Vesicles to quantify the mechanics of lipid membranes (bending and stretching moduli). (a) SEM image of a sliding element with a design adapted to GUVs, including 7 pipettes (12 µm in diameter), and fluorescence microscopy micrograph of 7 DOPC GUVs trapped within such pipettes inserted in a PDMS channel. (b) Fluorescence micrographs of a GUV blocked inside a 12-µm-diameter pipette, for three values of the pressure difference applied to the vesicle. Reference situation *ΔP*_0_, and two successive equilibrium position. The quasistatic increase of pressure causes a progressive increase of the GUV area, quantified from the length of the GUV protrusion within the pipette. (c) Evolution of the relative area increase as function of the tension for six DOPC GUVs of the same experimental run, and fitted curve according to equation (1). The displayed numbers correspond to the outcomes of the fitting for this particular experiment, as a typical example of data dispersion.

Integration was made by inserting the SMP in the PDMS chip. This step could be either achieved manually under a binocular microscope, with alignment precision between the pipettes and the fluidic channel in the order of 50 µm, or aided by a specific 3D-printed holder if better alignment was required (see Figure SI-2). In order to reduce friction, an anti-adhesive coating (fluorinated silane deposited in the gas phase) was realized on the SMPs before insertion. Insertion was also facilitated thanks to isopropanol lubrication, eliminated afterwards by evaporation (see Supplementary Information).

Once inserted, the SMP blocked the fluid in the main channel by letting it flow only through its cylindrical holes. The height and width of the guide were 20% smaller than the height and width of the sliding element it received (typically 450 µm for the guide and 550 µm for the sliding element), which we found optimum for elastic deformation to ensure a good sealing upon insertion. We checked the absence of leakage in the whole range of the pressure controller (325 mbar). Thus, when a vesicle or a spheroid was injected in the inlet solution, it was carried by the flow until it arrived in front of one of the micropipettes into which it was blocked and aspirated. In the design of the SMPs, the center of the pipette was placed at a Z position permitting objects to be trapped without touching the bottom of the fluidic channel, while being in focus under the microscope.

Chip operation differed slightly for GUVs and spheroids and are detailed in the next sections and in the Supplementary Information. Fluidic protocols shared some characteristics: after degassing and injection of the buffer to pre-wet the whole chip, the solution containing the objects of interest was injected to trap them at the pipettes. Measurements of the mechanical properties were achieved by quantifying deformations (of the GUVs or spheroids) under a programmed pressure sequence, by optical microscopy and image analysis.

This fabrication approach permitted the integration of cylindrical holes (or any extruded shape of arbitrary cross section) aligned with the main axis of fluidic channels, thanks to photolithography of two elements along two orthogonal planes, which can hardly be achieved by standard manufacturing techniques. This feature makes SIMPA technology uniquely suited for high throughput micropipette aspiration, which we demonstrate in the following sections.

### Micropipettes for vesicles: elastic moduli of lipid membranes

#### Background: membrane bending and stretching moduli, standard micropipettes

The mechanics of a lipid bilayer can be described by two main parameters: its resistance to bending, quantified by the bending modulus κ_c_; and its resistance to an increase of area per molecule (stretching modulus *K*_*A*_). These moduli determine how the area of a vesicle *A* increases with its tension *σ*, with reference to a state at low tension *A*_0_, *σ*_0_. The relative area increase, *α* = (*A* − *A*_0_)/*A*_0_, reads ^9,16^:

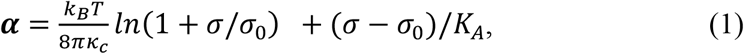

where *k*_*B*_ is the Boltzmann constant, and *T* the temperature.

The increase of area at low tension is mostly controlled by the smoothing of thermal fluctuations against bending (first term of equation (1)), whereas for a higher tension (typically 1 mN/m), it is set by the stretching modulus *K*_*A*_ (second term of equation (1)).

In standard micropipette experiments, a progressively increasing tension is induced thanks to a pressure difference *ΔP* applied to the vesicle aspirated in the pipette. With the hypothesis that the pressure inside the vesicle is equilibrated, and that the tension is homogeneous, the vesicle tension can be deduced from Laplace law according to:

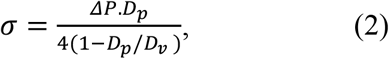

where *D*_*p*_ and *D*_*v*_ are the pipette and vesicle diameters, respectively. With the additional hypothesis of constant vesicle volume during the experiment (low permeability of the lipid bilayer to solvent, few minutes experiments duration), and a first order approximation 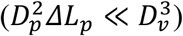, the area increase is deduced from *ΔL*_p_, the position of the vesicle protrusion within the pipette with respect to the reference state (*A*_0_, *σ*_0_), see Figure 2(b):

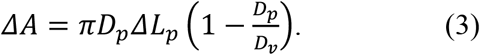

#### Chip operation and measurement protocols

These principles apply to our microfluidic chips: we designed channels (100 µm deep, 400 µm wide) in which fluorescently labelled GUVs with a typical diameter of 50 µm, obtained by standard electroformation (see Supplementary Information), could flow. The channels integrated sliding elements with up to 7 pipettes of diameter *D*_*p*_ ≃ 12 µm, see Figure 2(a). Since the hydraulic resistance of these pipettes were much larger than those of the inlet and outlet channels, the pressure drop applied to vesicles, *ΔP* in equation (2), was almost equal to the pressure drop applied on the whole channel *ΔP*_*channel*_, even in the case where some pipettes were not blocked by GUVs. After pre-treatment of the chip by a casein aqueous solution to avoid GUVs adhesion on the pipette walls, an aqueous solution of the fluorescently labelled GUVs electroformed in sucrose (see Supplementary Information) was gently injected within the chip (pressure controller, ΔP_channel_∼1mbar). Trapped GUVs within the pipettes were first prestressed at *ΔP*∼3 mbar (corresponding to a tension *σ*∼1 mN/m) for a few minutes to remove their possible defects. Then, the inlet pressure was set down to the value cancelling *ΔP* (GUV starting to escape the trap upstream). From this reference, the pressure was slowly increased by steps (3 s duration), in order to quantify the increase of *L*_*p*_ with *ΔP*, see Figure 2(b). The first step leading to a measurable GUV deformation was used as the reference state (*ΔP*_0_, *ΔL*_*p*_ = 0, *A*_0_, *σ*_0_), see top panel in Figure 2(b). Step height was increased from 0.01 mbar at the beginning (bending regime) to 0.5 mbar for the stretching regime. Fluorescence microscopy was used to image the GUVs. Being photosensitive, the dry films constituting the SMPs were slightly fluorescent, in particular for excitation wavelengths around 375 nm, as can be seen in the fluorescence characterization, see Figure SI-3. We chose fluorophores excited at higher wavelengths: excitation source at *λ*_*exe*_ = 541 nm, and emission filter at *λ*_*em*_ = 610 nm, which led to a fluorescence signal of the vesicles much stronger than the one of the dry film, see Figure 2(a). For large enough GUVs (*D*_*v*_ ≥ 2.5*D*_*p*_), standard image analysis was used to deduce *ΔL*_*p*_ as a function of *ΔP*. The relative area increase as a function of the tension was then calculated from Equations (2)-(3) for each GUV. The values of the bending and stretching moduli were then deduced by fitting Equation (1) to the experimental curve. As exemplified in Figure 2c showing six measurements realized in parallel, we have used a three-parameter fit, by letting the reference tension as a free parameter, in addition to the determination of *κ*_*c*_ and *K*_*A*_. It was found to more accurately reproduce the data trend in the bending regime than a two-parameter fit and fixed experimental reference tension *σ*_0*exp*_. The associated difference between *σ*_0*exp*_ and the fitted value was in the range *δσ*_0_ ≤ 10^−5^ mN/m, corresponding to a pressure difference *δP* ≤ 3 Pa. We independently characterized the accuracy on pressure control to be better than 0.5 Pa, so this value is a little higher than expected. We attribute this slight discrepancy to a higher uncertainty in determining the absolute value of the pressure, even though relative variations are precisely measured. With this procedure, the curve superimposed on experimental data both for bending and stretching regime, with a coefficient of determination of the fitting R^2^ ≥ 0.99.

#### Results for simple composition, effects of sugar and cholesterol

The results obtained with GUVs of simple composition (bilayer of the mono-unsatured lipids 1,2-dioleoyl-sn-glycero-3-phosphocholine, DOPC in sucrose solutions) are summarized in the histograms of Figure 3(a). The statistics is slightly lower for the stretching modulus *K*_*A*_ (N = 41) than for the bending modulus *κ*_*c*_ (N = 59) because some GUVs escaped the pipettes at moderate pressure, without fully entering the stretching regime. We deduced the value of *K*_*A*_ only for vesicles escaping at a tension *σ* ≥ 0.75 mN/m. This fragility, which can be attributed to dispersion in the lysis tension, possibly due to minor defects in some GUVs, was not correlated to the measured value of *K*_*A*_ and *κ*_*c*_.

**Figure 3.**
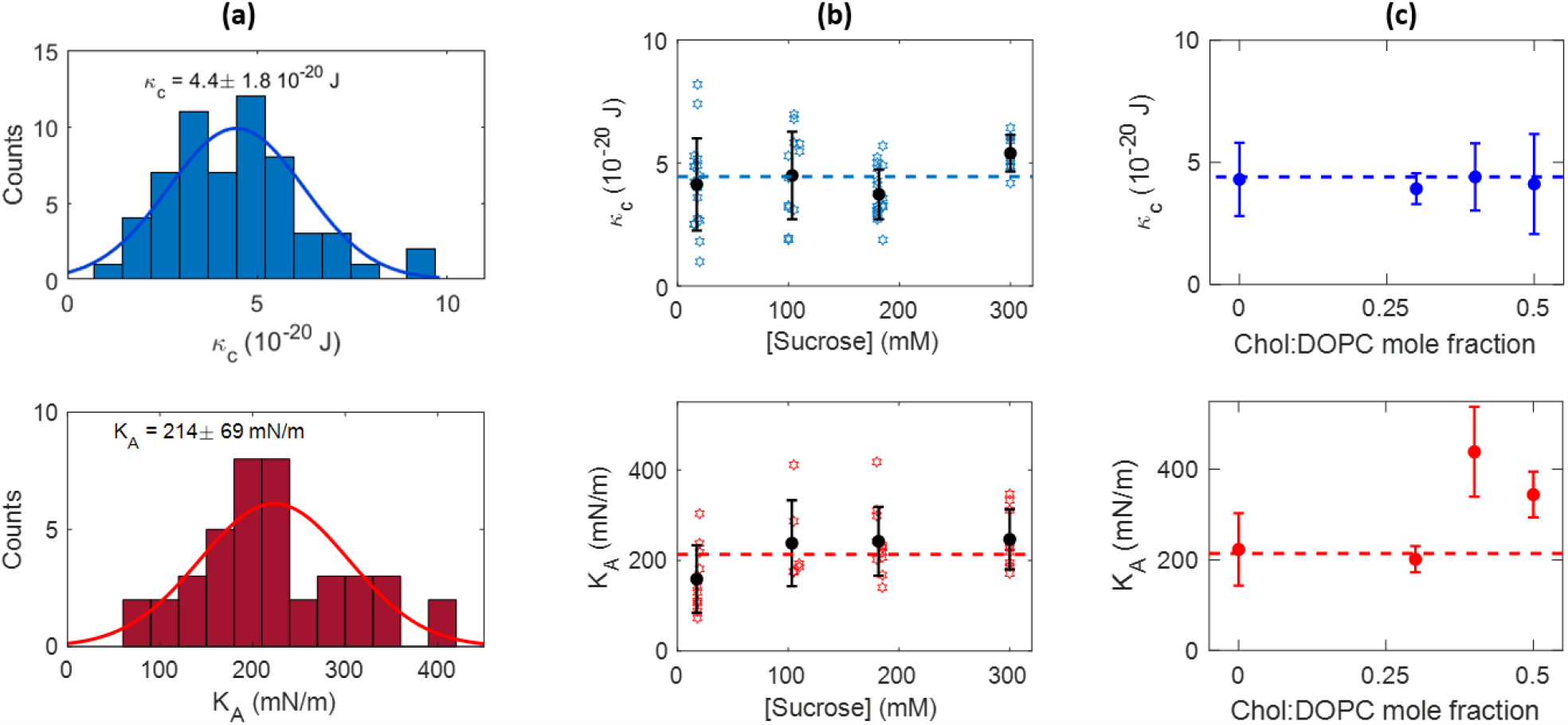
Bending and stretching moduli of lipid bilayers. (a) Histograms of the bending (top) and stretching (bottom) moduli of DOPC membranes, and associated Gaussian fits. (b) Influence of the sucrose concentration on the value of the bending (top) and stretching (bottom) moduli, for DOPC membranes. (c) Influence of cholesterol on the bending (top) and stretching (bottom) moduli of mixed DOPC-cholesterol vesicles, as a function of the cholesterol:DOPC mole fraction.

We also investigated the effect of the sucrose concentration on the bilayer mechanics, for DOPC lipids, see Figure 3(b). No systematic variation of both bending and stretching moduli were observed from 15mM to 300 mM, within our experimental error.

Finally, we performed measurements on bilayers composed of DOPC mixed with up to 50% cholesterol, Figure 3(c). We observed no dependence of the bending modulus on the cholesterol/lipid molar fraction, whereas the stretching modulus almost doubled for molar fraction 0.4 and 0.5.

#### Discussion: on-chip pipette to characterize vesicles mechanics

Overall, the results in Figure 2 and Figure 3 show that the proposed approach is suited to determine the mechanical properties of lipid membranes, similarly to the classical micropipette aspiration. However, the throughput of our method is higher (roughly multiplied by the number of pipettes in parallel, 7 in Figure 2) since several GUVs can be characterized in parallel. The integration in a microfluidic device has also the advantage of avoiding the manual, tedious search of vesicle and micromanipulation of the pipette, since the driving flow in the channel naturally brings the vesicles to the pipettes, and facilitates the trapping of GUVs. In addition, pressure controllers used in routine for microfluidic setups have sub-second response time and permit the automation of pressure vs. time protocols.

The obtained values are reasonably consistent with the literature. For DOPC at room temperature, the determined stretching modulus *K*_*A*_ = 214 ± 69 mN/m falls within the range of most micropipette measurements (respectively *K*_*A*_ = 210 ± 25 mN/m, 198 *m*N/m, and 265 ± 18 mN/m for references ^30–32^). The bending modulus we obtained (*κ*_*c*_ = 4,4 ± 1.8 .10^−20^ J) is in the lower limit of published values for measurements with micropipettes (respectively *κ*_*c*_∼9.1 ± 1.5 .10^−20^ J, 8.5 10^−20^ J, 4.7 10^−20^ J for DOPC in references ^30,32,33^), reported in a recent review ^34^ to be in the range *κ*_*c*_ = 4 − 16 10^−20^ J for monounsaturated lipids. It has to be mentioned that systematic differences between groups and measurement method are thoroughly discussed, and only partly explained by differences in the probes scales or experimental protocols, in several reviews^9,34–36^.

The dispersion of our data is a bit higher than in the literature (coefficients of variation 41% and 29% for *κ*_*c*_ and *K*_*A*_ respectively). GUVs synthesized via electroformation have inherent variability. We also attribute the dispersion to the fact that the only eliminated GUVs were those with diameter *D*_*v*_ ≤ 2.5*D*_*p*_), or with visible defects (such as internal vesicles), contrary to standard micropipette aspiration where the operator arbitrarily chooses the GUV to be probed. The absence of influence of sugar concentration we observed (Figure 3(b)) is consistent with most recent observations and discussions of the literature, even though this is still a quite controversial issue.^2,36–38^

When varying the membrane composition by mixing DOPC with cholesterol, we observed no change in the bending modulus, from pure DOPC up to the maximum cholesterol content tested (0.5mol/mol). On the opposite, a two-fold increase was observed for *K*_*A*_ for increasing cholesterol content, with a possible threshold between 0.3 and 0.4 molar fraction in cholesterol. These observations are completing a rich literature on the issue of cholesterol influence on membrane structure and properties. Bending rigidity was shown to be strongly lipid dependent^39,40^, stiffening by cholesterol being observed only for saturated lipids, with no effect for mono-unsatured lipids such as the DOPC used in the present study^41,42^.

Finally, since microfluidic lateral channel can be integrated to the design, the approach is well-suited for temporal change of the chemical environment surrounding the GUVs. It opens interesting perspectives to investigate for example the kinetics of interaction of lipid bilayers with biomolecules or relevant synthetic entities (molecules, macromolecules or nanosystems).

### Micropipettes on cell aggregates: quantifying spheroids’ rheology

#### Background: Viscoelastic model for biological tissues, standard micropipettes

Many biological tissues behave as viscoelastic fluids, which is both due to the properties of individual cells (cytoskeleton, nucleus), and to the way they assemble in the tissue (extracellular matrix, adhesion between cells). Thus, when a spheroid (simple 3D cell aggregate) is probed by micropipette aspiration with a pressure step, it reacts with two different regimes. First, an instantaneous deformation is observed, directly linked to the tissue’s elastic properties. Then, over time, the tissue flows into the micropipette like a viscous fluid. Several viscoelastic models describe this type of material, but the modified Kelvin-Voigt shown in the insert of Figure 4b is the simplest that closely reproduces the response observed in Figure 4b. It consists of a Kelvin-Voigt element (spring *k*_1_ in parallel with damper µ_c_), modified by the spring *k*_2_ to account for an instantaneous elastic response, in series with a dashpot µ_t_, which corresponds to long-term viscous flow. In this description of the tissue as a soft material, viscosity and elasticity are completed by the aggregate’s surface tension *γ*, excess of surface energy that originates from a combination of the interaction between cells and differences in cortical tension between the peripheric and the core cells^12,43^. In a standard micropipette experiment, a spheroid of radius *R* is aspirated in a pipette of radius *R*_*p*_ with a suction pressure Δ*P*. The effective force inducing spheroid deformation reads: 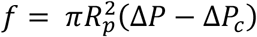, with 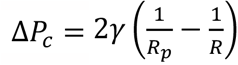 the Laplace pressure generated by the curvature imposed by the pipette. Δ*P*_*c*_ corresponds to the minimum pressure needed for the spheroid to continuously flow inside the pipette. For Δ*P* > Δ*P*_*c*_, the spheroid’s response to a pressure step can be written, in terms of its temporal elongation *L*(*t*) inside the pipette (see Figure 4(a)):

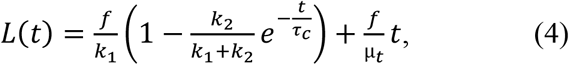

where 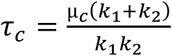 is a viscoelastic characteristic time.

**Figure 4.**
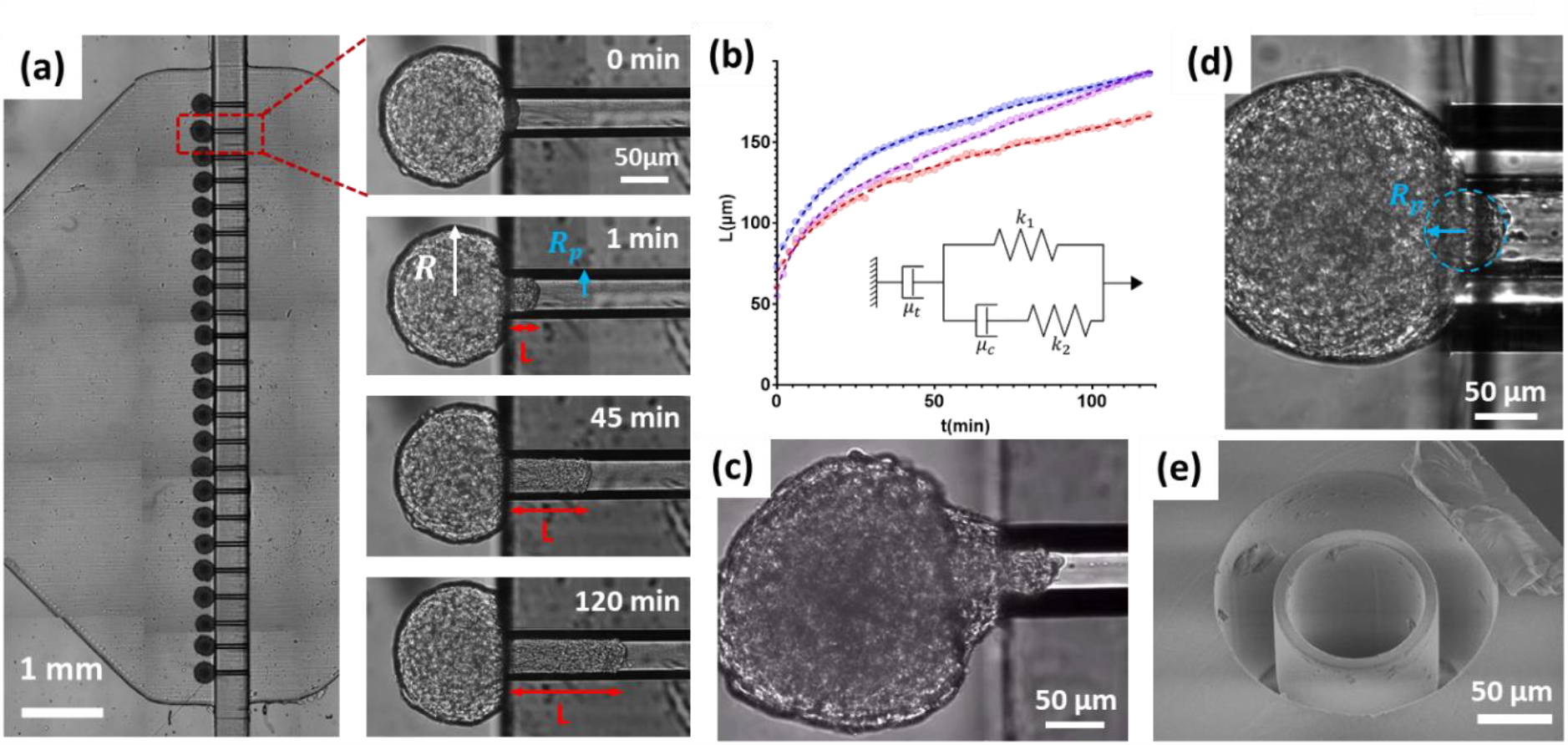
On-chip pipettes applied to 3D cellular aggregates to quantify their viscosity and elasticity. (a) Micrograph of 23 spheroids trapped in the pipettes in the SIMPA chip. Close-up on a single micropipette: time lapse of the aspiration of one A338 spheroid submitted to a pressure step *ΔP* = 50 mbar from t = 0 s. (b) Evolution of the spheroids’ positions *L*(*t*) in the pipette as function of time for 3 simultaneous parallel measurements, and fitted curves according to equation (4). (c) Micrograph of a spheroid just after release of the aspiration pressure. The conical shape indicates that Laplace pressure is not the only process expelling the spheroid from the pipette, during retraction. (d) Micrograph of a spheroid aspirated with a pressure just equal to the Laplace pressure *ΔP*_*c*_, leading to a radius equal to the pipette’s radius. (e) SEM image of the pipette design with thin wall, used to improve the optical quality of the image in (d).

The first term in equation (4) refers to a viscoelastic solid, with two elastic moduli acting at two timescales: a first modulus *E*_*i*_ = (*k*_1_ + *k*_2_)/*πR*_*p*_, associated with an instantaneous deformation of the spheroid, and a second elastic modulus *E* = *k*_1_/*πR*_*p*_, which comes into play after a typical time *τ*_*c*_. These two elastic moduli are usually attributed to the cellular cytoskeleton’s reaction to pressure: the elasticity of the actin cortex is first assessed, fibers then rearrange, leading to a softer long-time elastic response.

The second term describes flow at the tissue level and it corresponds to the constant speed flow of a fluid of viscosity *η* = *μ*_*t*_/3*π*^2^*R*_*p*_ inside the pipette, with the hypothesis that viscous dissipation occurs only at the inlet, due to cell rearrangements. As detailed in reference ^18^, this regime neglects wall friction, which is achieved thanks to surface treatment limiting cell adhesion on the pipette’s walls.

#### Chip operation and measurement protocols

We have developed microfluidic chips enabling parallel aspiration of spheroids, see Figure 4(a). The major part of spheroid experiments was conducted on A338 mouse pancreatic cancer cells containing a KRas^G12D^ mutation. We also realized measurements on a S180 murine sarcoma cell line. Spheroids were cultured for 48 to 72 hours either using the suspended droplet technique, or in an array of agarose wells obtained from a 3D-printed mold (see Supplementary Information).

Regarding fluidic design, most results shown in this paper were obtained with a design probing 5 spheroids in parallel, but we have demonstrated the aspiration of up to 23 spheroids, see Figure 4(a). Channel height was 450 µm to accommodate all spheroid sizes. The chamber width was 10 mm for the 23-position chip (2 mm for the 5-position chip). A single microfabrication run allowed us to manufacture around 150 SMPs, see photo in Figure 1(b), which permitted us to test different pipette diameters and designs. Most experiments were conducted with 70 µm diameter pipettes, chosen as a compromise: much smaller than spheroids size, and significantly larger than cells size, for the granular nature of the tissue not to be too critical for the continuous description of the rheological model. Pipette were 500 µm in length.

The experimental protocol is detailed in the Supplementary Information. Briefly, after insertion of the SMP, the chip was placed under vacuum to eliminate residues of isopropanol and to limit bubble formation during experiments. The chip was then prewet by the culture medium under the microscope in an environmental chamber (5% CO_2_, 37°C). The spheroids were gently aspirated within a PTFE tubing, then they were loaded into the chips by applying Δ*P*_*load*_∼1 mbar. With this low pressure difference, the spheroids were transported to the pipettes under limited stress. They also hardly deform and did not start flowing continuously inside the pipettes since Δ*P*_*load*_ is significantly lower than the critical pressure ΔP_c_. The microscope stage and the camera were configured to record timelapse sequences of the aspiration of all the (5 or 23) trapped spheroids, with typical time interval 30 s, total experiment duration 3 hours. A pressure step, synchronized with image acquisition, was then applied with Δ*P* typically set to 50 mbar.

In order to validate quantitatively the SIMPA approach with respect to MPA, we performed measurements on the murine sarcoma cell line S180-GFP that was characterized by Guevorkian *et al*.^17^ by MPA. Since Laplace pressure contributes to the spheroid’s flow (see Equation (4)), the surface tension *γ* needs to be determined in order to deduce the viscoelastic parameters. Like in reference ^17^, aspiration was followed by retraction experiments, in which Laplace pressure is the only source of movement. The histograms of measured viscoelastic parameters are plotted in Figure SI-4 (N = 23). We obtained *γ* = 10.8 ± 2.4 mN/m, η = 1.37 ± 0.03 10^5^ Pa. s, E = 213 ± 17 Pa, *E*_*i*_ = 773 ± 47 Pa. The values of the viscosity and long-time elasticity are fully consistent with the results reported in reference ^17^ (η = 1.9 ± 0.3 10^5^ Pa. s, elastic modulus deduced from an average of relaxation times: E = 700 ± 100 Pa), given that the cell line may have slightly evolved since 2010, and more importantly that the culture conditions to form the spheroids were not exactly the same in the two studies.

For A338 mouse pancreatic cancer cells spheroids, a typical timelapse for one position is shown on the right panel of Figure 4(a), and in Supplementary Video 1. The position of the spheroid protrusion as a function of time *L*(*t*) was determined by a custom image segmentation algorithm described in Supplementary Information, see Figure SI-5 and Supplementary Video 3. Typical results of a single experiment driven on A338 spheroids are displayed in Figure 4(b), together with the fit of these results by equation (4).

However, regarding surface tension, we observed for this cell line a complex conical shape of the spheroid upon retraction, see Figure 4(c) and Supplementary Video 2, and its fast ejection from the pipette. Several mechanisms could explain this behavior: stored elastic energy could contribute to expelling the spheroid out of the pipette (similarly to what is mentioned in reference ^22^); and additionally thanks to low wall friction the spheroid could slide upstream without dissipation and progressively round up at the pipette’s corner because of surface tension. We thus used alternatives to such retraction experiments and measured *γ* by directly characterizing Laplace pressure thanks to other sets of experiments. We quantified the minimum critical pressure leading to continuous flow of the spheroid Δ*P*_*crit*_, which should also correspond to the pressure for which the radius of the spheroid meniscus (formed by cells at its surface within the pipette) equals the pipette radius, see Figure 4(d). This set of experiments was realized on a specially designed thin-wall pipette, see Figure 4(e), to improve the quality of optics. Both pressures were determined to be very close and equal to Δ*P*_*c*−*crit*_ = 5 ± 0.5 mbar). These measurements led to a value of the surface tension *γ*_*crit*_ = 10 ± 1 mN/m. We also quantified the ratio *γ*/η from the dynamics of spheroid fusion^44^, see Supplementary Information, Figure SI-6. These independent off-chip experiments led to *γ*_*fusion*_ = 4.5 ± 0.9 mN/m. The fusion experiment mainly probes the external layers of the spheroid, and surface tension of cell aggregates was recently discussed theoretically to be a multi-scale complex concept^45^, so that different configurations could lead to slightly different results. The viscosity retained for this fitting was the one deduced from aspiration experiments. In addition, note that in both cases, the value corresponds to the surface tension at low stress, referred to as *γ*_0_ in previous studies which have evidenced a possible increase of *γ* upon aspiration^17^. We finally retained the on-chip measured value *γ*_*crit*_, since it was determined in the same flow configuration as the pipette aspiration. It is in the typical range of literature measurement of biological tissues’ surface tension^46^, even though most available data are on less cohesive configuration than the epithelial one probed here. Let us mention that since the applied pressure in aspiration experiments was significantly higher than the typical Laplace pressure, an error on surface tension determination would not critically affect the determination of viscoelastic parameters.

With this value of the surface tension, we extracted the rheological parameters from the fitting of the experimental curves *L*(*t*) with equation (4), see Figure 4(b). The fittings closely reproduced the trends of the experiments. Graphically, tissue viscosity is deduced from the slope at long time, whereas the first (short-time) elastic modulus *E*_*i*_ can be deduced from the initial instantaneous elongation of the spheroid, the second modulus *E* from the intercept of the long-time linear flow regime with the vertical axis, and *τ*_*c*_ from the typical time scale to reach this regime.

#### Measuring viscoelastic properties of spheroids: results and discussion

The results obtained with A338 spheroids are shown in Figure 5(a). All (*N* = 134) measurements were realized with the 5-pipettes design, in about 30 experiments, each lasting a few hours, which demonstrates the high throughput of the method. We measured elastic moduli *E* = 1.4 ± 0.5 kPa, *E*_*i*_ = 2.5 ± 0.9 kPa, and a time scale *τ*_*c*_ = 15.3 ± 8.1 min (error bars indicates the standard deviation). For viscosity, the distribution was observed to be better fitted by a log-normal distribution than by a Gaussian. The maximum (mode) of the fitted distribution was *η*_*ln*_ = 1.20 MPa. s, with a distribution width *σ*_*η*−*ln*_ = 0,67 MPa. s. We observed a more reduced dispersion between spheroids of the same batch: the average of standard deviations deduced from single experiments (5 or 23 simultaneous measurements) was *σ*_*η*−*batch*_ = 0.5 MPa. s and 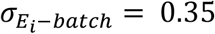 kPa for the viscosity and short-time elasticity *E*_*i*_ respectively. These values can be interpreted as an upper bound of the measurement uncertainty, demonstrating the reproducibility of the technique. The width of the histograms in Figure 5(a) mostly originates from biological variability.

**Figure 5.**
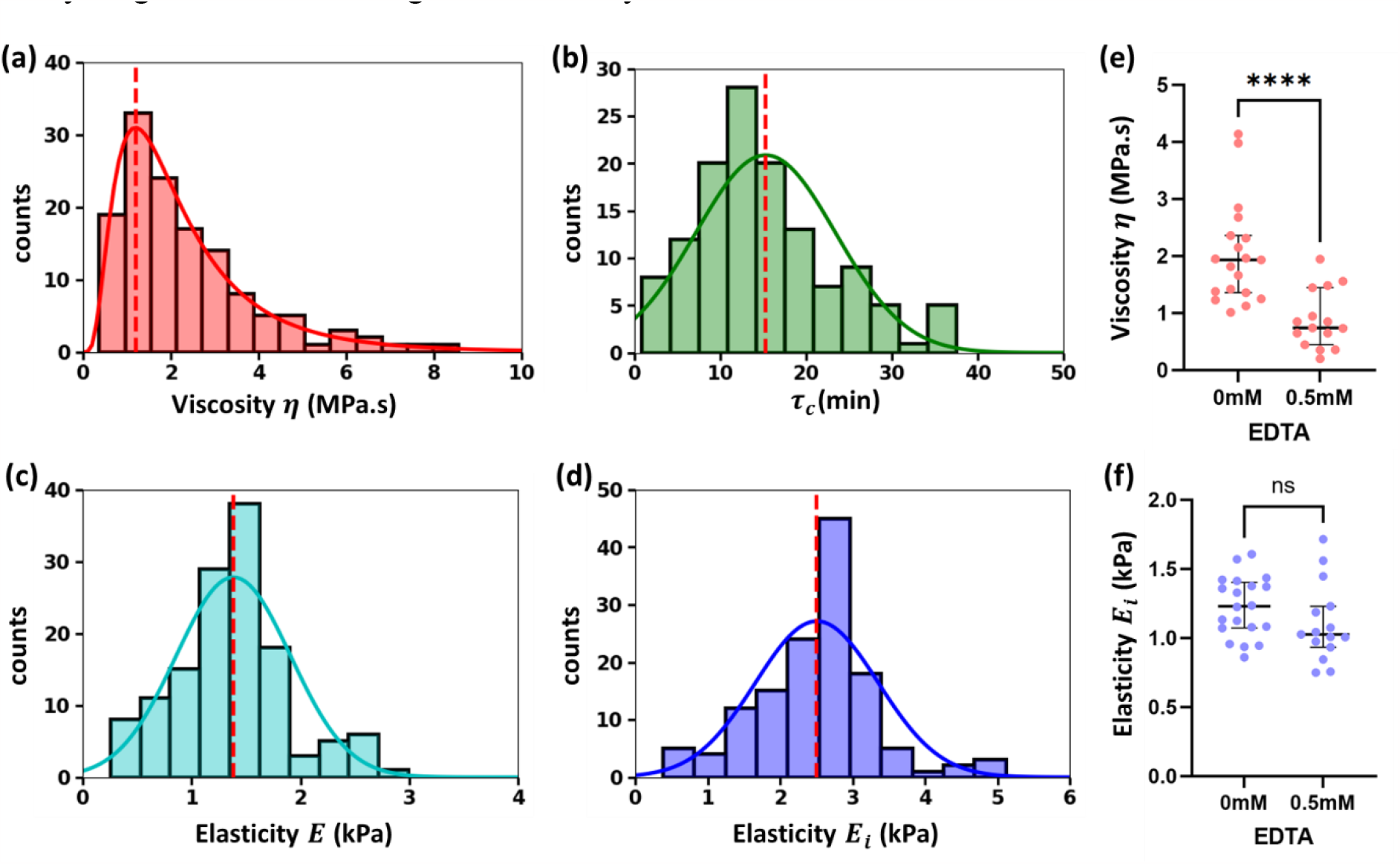
Viscoelastic properties of A338 cellular aggregates: histograms of the viscosity (a), characteristic viscoelastic time (b), long-time elasticity (c), short-time elasticity (d). (e-f) Influence of EDTA on the spheroids’ viscosity and short-time elasticity.

To the best of our knowledge, no viscoelastic measurements have been published for this cell line. However, the high value of viscosity and elasticity, about ten times the typical values measured for very dynamic embryonic tissues^47^, are consistent with the strong cohesion of pancreatic epithelial-like tissues.

We also assessed the effect of Ethylenediaminetetraacetic acid (EDTA). EDTA affects adhesion between cells by chelating metallic ions, including calcium, necessary for adhesion proteins to operate. We incubated the cells with EDTA during formation of the spheroids before measuring viscoelastic properties. The results are shown in Figure 5(e-f). The viscosity was significantly reduced for spheroids incubated with EDTA with respect to the control, whereas short-time elasticity was not affected. This behavior is consistent with a reduced adhesion facilitating rearrangement of cells (T4 events^25^), leading to decreased viscosity, whereas elasticity, originating mostly from cells’ cytoskeletons, was not strongly impacted.

We now discuss the specificities of the SIMPA technology for spheroid rheology, with respect to existing methods.

First, the approach benefits from the advantages of standard MPA: it is quantitative, it probes optically-determined locations of an object, which opens the possibility to test different zones of a tissue for non-spherical aggregate. MPA applies forces from the external cell layers, which can give complementary information to methods applying homogeneous stress, like magnetic rheometry^15^. In our chips, since it is the microfluidic flow that pushes the spheroids towards the pipettes, spheroid’s orientation and the precise point they contact the pipette’s inlet cannot be controlled by the operator independently of the fluidic design, which can appear as a limitation. However, for non-spherical objects it could turn into an advantage: the shape of the upstream channel and the location of pipettes on the sliding element could be specifically designed to set this orientation and probe well-defined areas.

The SIMPA technology has unique features compared to standard MPA: the throughput is multiplied by the number of spheroids that can be probed in parallel (demonstrated to be up to 23 in this article). In addition, the chip format permits the use of low volumes of sample (typically a few hundred µL), with an easy spheroid loading since the flow naturally pushes the spheroids to the free SMPs. The chips can also be washed and reused, and the spheroids extracted out of the chip for further characterization. It is possible to keep spheroids for long times (we observed spheroids stable for three days with no visible necrosis). This comes from the environmental chamber surrounding the chip on the microscope (temperature set to 37°C and 5% CO_2_), but also from PDMS permeability to oxygen, and from a fast diffusion of nutrients within the chips.

## Conclusion

We present in this paper the SIMPA technology, a parallel, quantitative integrated aspiration micropipette method. We demonstrate its relevance to characterize quantitatively mechanics both at the cell membrane scale and at the multicellular scale. With respect to standard MPA, its throughput is multiplied by the number of pipettes in parallel, shown to be for this proof of concept 7 and 23 for GUVs and spheroids respectively. With respect to other integrated on-chip micropipettes^22,24,25^, our approach is the only one that combines circular geometry and parallel probing, in a user-friendly format. Thus, even if interesting analyses have been developed recently for squares or rectangles^23^, circular traps are quantitative by design, they fully eliminate both anisotropy of the constraints and residual flows in the corners.

Several perspectives emerge from the versatility of the method, related to fluidic design. As the most obvious evolution, larger fluidic chambers, or pipettes placed at different z-positions, could lead to an even larger throughput by adding further parallel pipettes, if required. More interestingly, changing the chip design permits controlling physico-chemical stimuli around trapped objects. First, pipettes surrounded by small holes, or including slits to let a fraction of the flow pass around trapped objects, could be used to probe vesicles or cells aggregates while submitted to a shear stress. By using cross-shaped pipettes, we have observed that shear stress affects lipid domains, as demonstrated in reference ^48^, see Figure SI-7. Secondly, adding lateral fluidic channels close to the sliding element permits changing in real time the chemical environment of trapped GUVs or spheroids, see Figure SI-8. As a proof of concept, we demonstrated the dynamic exposure of trapped spheroids to microparticles, see Supplementary Videos 4 and 5. This type of design could be relevant to study the response to drugs at short timescales, typically seconds or minutes, or to apply spatio-temporal stimulations such as the ones originating from heart beating or circadian cycle. Quantifying the influence of different drugs, at different timescale, should improve our understanding of the microscopic origin of tissue rheology. In the same perspective, the technology can apply a dynamic pressure stimulus, as in reference ^15^, which is a relevant way to assess the validity of different rheological models.

As a further perspective, we propose to extend the single vesicle configuration to the probing of single cells. For that purpose, since typical mammalian cells are 6-10 µm in dimension, pipettes with diameters of order 2-5 µm are required. We were able to block and release A338 individual cells in suspension in 4 µm-diameter pipettes (see Supplementary Video 6).

Finally, specific versions of the technology can be developed to improve the quality of optics (thinner walls, see Figure 4(c), or glass versions of the sliding elements, see Supplementary Video 7). Overall, the SIMPA technology will help identify how collective properties emerge from individual cell deformations and rearrangements.

## Supporting information

Supplementary Info and Legends for Supplementary Videos

## Acknowledgements

This work was supported in part by CNRS through the PICS CNRS program “Microfluidics for Soft Matter”, and by INSIS-CNRS. This work was partly supported by LAAS-CNRS micro and nanotechnologies platform member of the French RENATECH network. Funding by “ADI, Université de Toulouse, Région Occitanie”, and by Federation Fermat, Université de Toulouse, are acknowledged. We thank Benjamin Reig for SEM imaging, Sandrine Souleille for microfluidics experiments, Charline Blatché for cell culture, Julien Roul for help in microscopy experiments. We thank Karine Guevorkian and Gregory Beaune for gently providing the S180 cell line.

## Author Contributions

ME, HA, MP realized experiments on GUVs. SL, ME, PL realized experiments on spheroids. SL, ME, PL, HA, MP, AL, FM, DB contributed to chips fabrications. LM, JB calibrated the photolithography process. DB, CM, MD, CR, PJ contributed to experiments and supervised the research. PJ and SL wrote a first draft of the manuscript, all authors read and amended the manuscript.

