## Supplementary Info and Legends for Supplementary Videos for "Parallel on-chip micropipettes enabling quantitative multiplexed characterization of vesicle mechanics and cell aggregates rheology"

### Microfabrication protocols: molds, sliding elements, PDMS chip

The fabrication of the molds for the PDMS chips and the sliding elements was carried out in a clean room, utilizing SUEX dry film photoresist sourced from DJ Microlaminates. Employed thicknesses were 50, 100, 250, and 500 µm. Some adjustments were made to the recommended microfabrication parameters provided by the manufacturer. The UV exposure was slightly reduced for thick films, to reach the high aspect ratio of the micropipettes. To ensure photoresist adhesion to the silicon wafer during mold formation, oxygen plasma was employed before lamination. Conversely, for the sliding element fabrication, resin detachment after development was achieved by utilizing silicon wafers coated with a ~100nm Cu/Ti layer. Subsequent to the fabrication process, this sacrificial layer was etched using an $NH_3/H_2O_2$ mixture. To prevent adhesion of PDMS to the molds and spheroids to the sliding element, a gas-phase treatment was realized using a Surface Preparation Deposition Memsstar machine. It consisted in grafting a self-assembled monolayer of Perfluorodecyltrichlorosilane (FDTS), after a deposition of a 10 nm-thick $SiO_2$ layer that ensured adequate grafting.

The PDMS chips were prepared by standard soft lithography, using a 1:10 ratio of Sylgard 184 PDMS, mixing the crosslinker with the base polymer. After crosslinking atop the molds, the formed PDMS was carefully unmolded and punched at the designated inlet and outlet points. A second, 50 µm-thick PDMS layer was obtained by spin-coating uncrosslinked PDMS onto a clean wafer at 400 rpm for 1 minute. The two PDMS part were bonded together by plasma.

### Preparation of Giant Unilamellar Vesicles:

Lipids were bought from Avanti Lipids. 1,2-dioleoyl-sn-glycero-3-phosphocholine (DOPC) lipid in powder was dissolved in chloroform at 0.5 mg/mL concentrations. The fluorescent dye Lissamine rhodamine B sulfonyl (1,2-dimyristoyl-sn-glycero-3-phosphoethanolamine-N-) dye was added at 0.1% molar ratio to the lipid solutions. Electroformation was realized according to standard protocols[1]: 10 µL of the lipid solution were added on two glass slides covered with ITO, which were placed under vacuum for two hours. The two glass slides were placed with the lipid deposits facing each other, spaced by a 1 mm thick O-ring. 200 µL of solutions of sucrose in water (concentration ranging from 15 mM to 300mM) were added before sealing. A 2 V peak-to-peak sinusoidal voltage at 10 Hz was applied for 3 hours on the film. The obtained GUVs were then diluted 5 times in a sucrose water solution of similar but 10% lower sucrose concentration, in order to prestretch the GUVs and reduce the number of membrane defects resulting from electroformation.

For cholesterol-DOPC mixtures, cholesterol was first dissolved in chloroform at 0.5 mg/mL concentrations. This solution was mixed to a 0.5 mg/mL DOPC solution in order to obtain 0.3, 0.4, and 0.5 molar ratio of cholesterol. The same fluorescent dye as for the DOPC experiments was used.

### Cell Culture and Spheroid Preparation Protocol:

For the majority of the experiments, we utilized a murine pancreatic cell line with the KRas$^{G12D}$ mutation (A338), characteristic of approximately 80% of pancreatic cancers. We also used a murine sarcoma fluorescent cell line (S180 GFP). The media used for the cell culture was Dulbecco's Modified Eagle Medium (DMEM), supplemented with 10% Fetal Bovine Serum (FBS) and 1% of an antibiotic solution (penicillin and streptomycin). Spheroids were generated either by the 20 µL hanging drop method following the protocol described in reference [2], or on structured 2% agarose cushions facilitated by a home-made 3D-printed pad. In both methodologies, seeding between 100 and 1000 cells per drop or well resulted in the formation of spherical aggregates within 48 hours. The resultant spheroids ranged from 100 to 300 µm in diameter, depending on the initial cell count.

### Microscopy and Fluidics Protocols:

Most experimental observations were conducted using a ZEISS Observer 7 microscope in bright field mode, employing a 20x air objective (NA = 0.8). The motorized stage was equipped with a TOKAIHIT incubation chamber, maintained at 37 °C with 5 % $CO_2$. The "Definite Focus" systems kept the focus during the experiments.

The microfluidic system comprised a 4-channel pressure controller (Fluigent MFCS), 0-69 mbar range and 0-25 mbar range for cell aggregates and GUVs respectively. Reservoirs were filled with media, and connected to 0.8 mm internal diameter plastic tubing (PTFE).

The PDMS chip was first rinsed with isopropanol, and the sliding element was also lubricated with isopropanol to ensure its good insertion. After insertion and 30 minutes of drying under vacuum, the chip was filled with the buffer (culture media for experiments on spheroids, sucrose solution for GUVs) from the outlet with a syringe; to clean the chamber and eliminate air bubbles thanks to PDMS permeability to gas. The outlet was then plugged to a reservoir of buffer connected to the pressure controller. Spheroids were first aspirated with a syringe in a tube, which was then inserted in the inlet of the chip and plugged to a reservoir of media connected to the pressure controller. For GUVs, the solution containing the vesicles was directly put in the reservoir connected to the pressure controller and to the chip inlet.

### Roughness of the pipettes

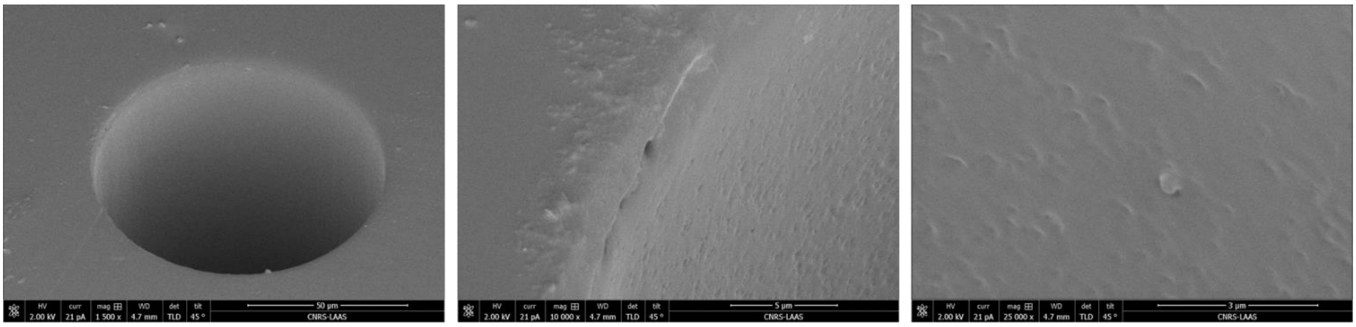

Figure SI-1- SEM micrographs of a 80-µm-diameter pipette, realized in SUEX dry film. From left to right: low, medium, high magnification, evidencing a very low roughness of the pipette. Typical defects are less than 20 nm in height.

### Controlled insertion of the SMP (Sliding Micro Pipettes)

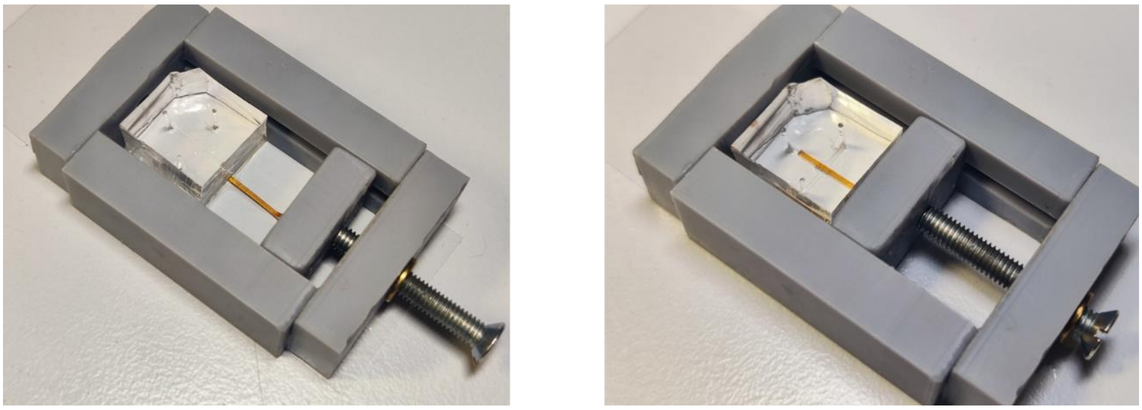

Figure SI-2- Holder enabling precise insertion of the SMP inside the chip. With this element, and under a binocular microscope, alignment of the pipettes with the fluidic channel is estimated to be 25 µm, instead of 100 µm for a manual insertion

### Fluorescence properties of the SMP fabricated in dry film

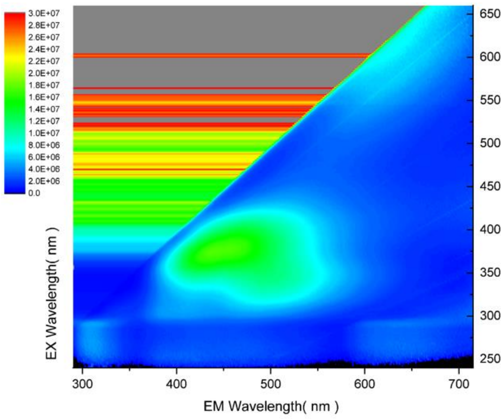


Figure SI-3 –Fluorescence spectroscopy characterization of the dry films (SUEX) SMPs (Arbitrary Units). A significant signal appeared for 400-520 nm emission wavelengths, when the SMP was illuminated with 340-430 nm wavelengths.

2D Fluorescence map acquired using a Fluorolog 3 fluorimeter from Horiba.

**Measurement of the viscoelastic properties of spheroids made with S180 murine sarcoma cell line**

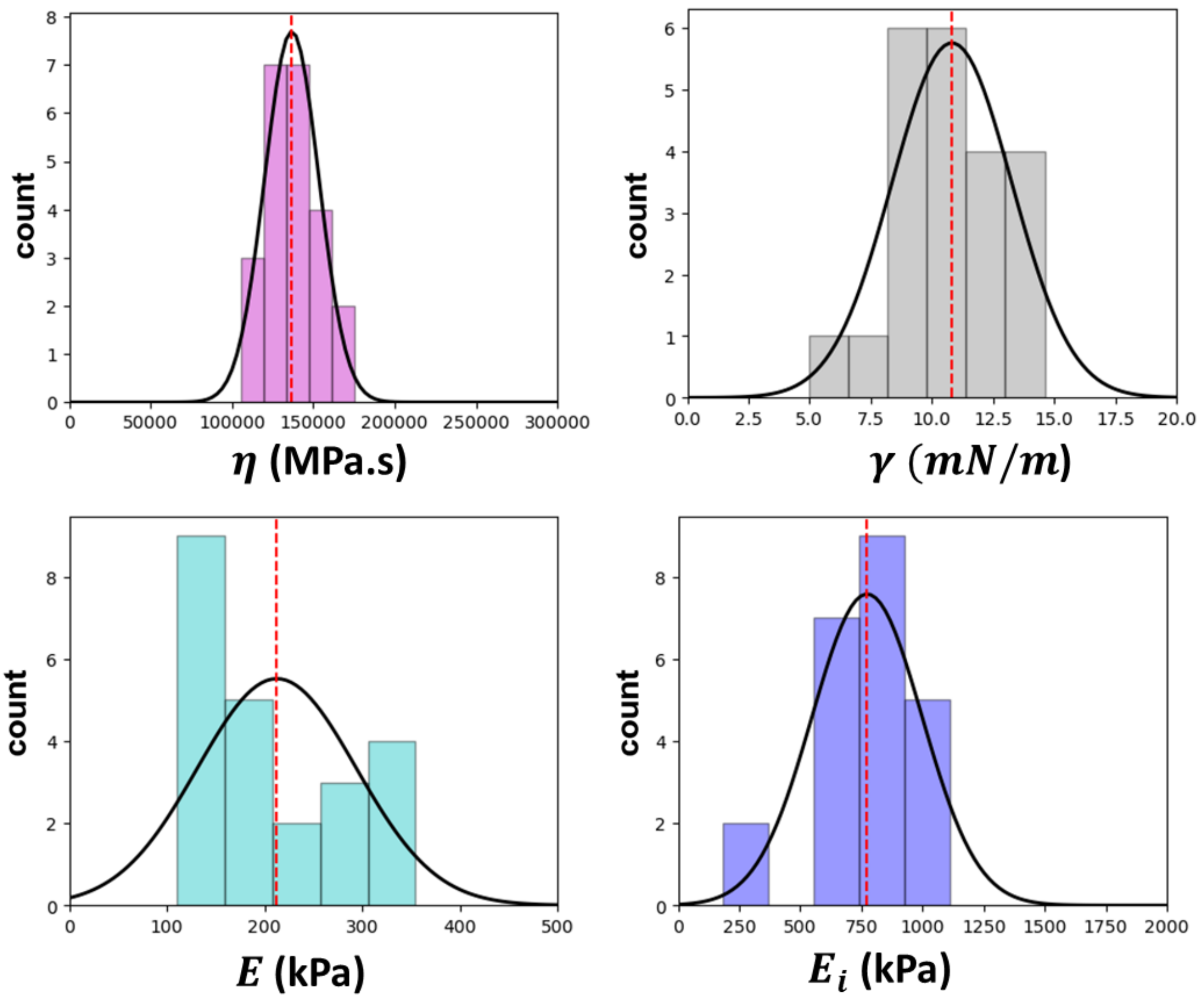


Figure SI-4- Measurement of the rheological properties of S180 murine sarcoma cell line. $\gamma = 10.8 \pm 2.4$ mN/m, $\eta = 1.37 \pm 0.03\ 10^5$Pa. s, E $= 213 \pm 17$ Pa, $E_i = 773 \pm 47$ Pa.

### Image analysis algorithm

To automate the identification of the spheroid intrusion in the pipette, we formulated an image analysis algorithm. The procedure begins with computing the image's derivative along the pipette axis. As illustrated in Figure SI-5, post-thresholding utilizing the Otsu method results in the vanishing of the pipette's edges, leaving predominantly the aspirated tongue in view. Subsequent dilation, erosion, and filling operations yield a predominant region representing the aspirated tongue, alongside minor regions attributable to image imperfections and debris. By quantifying the area of these segmented regions, we selectively retain the largest, which aligns closely with the tongue's contour. The tongue's tip is subsequently identified by fitting a circle to the contour points that represent the meniscus, ensuring we sidestep any distortions possibly caused by cells or vesicles disengaged from the tongue.

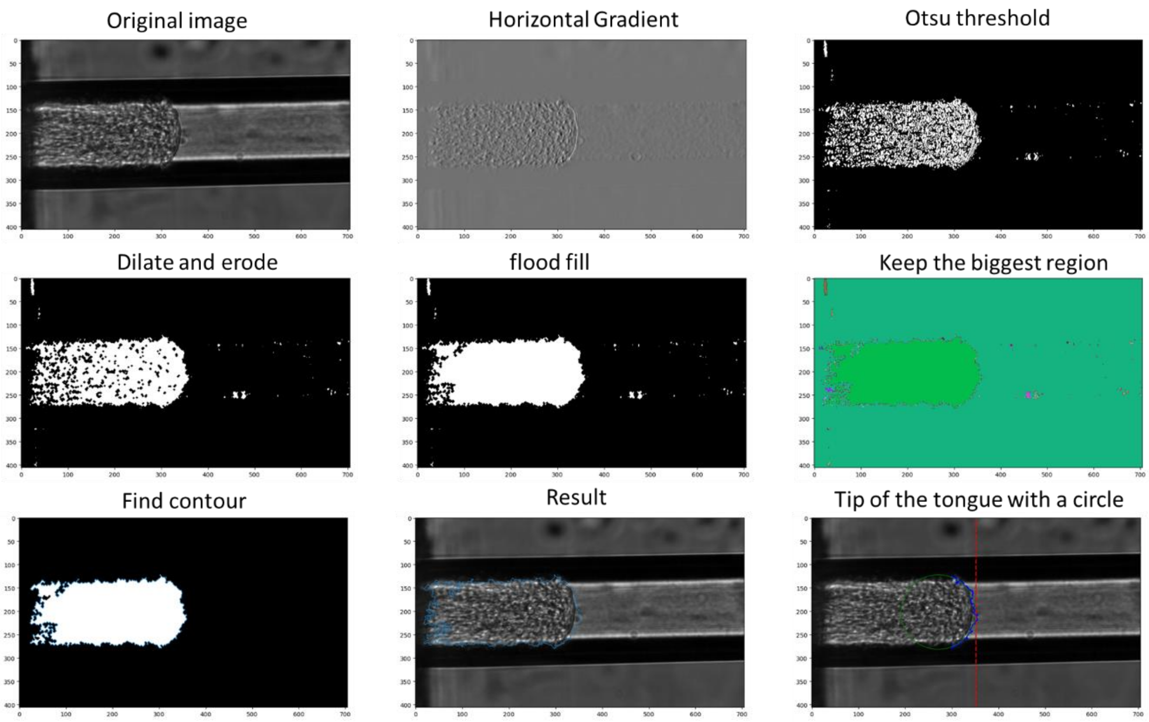


Figure SI-5 – Main steps of the image analysis algorithm, aiming at extracting the position of the spheroid's meniscus within the pipette.

### Independent off-chip measurement of the ratio surface tension / viscosity

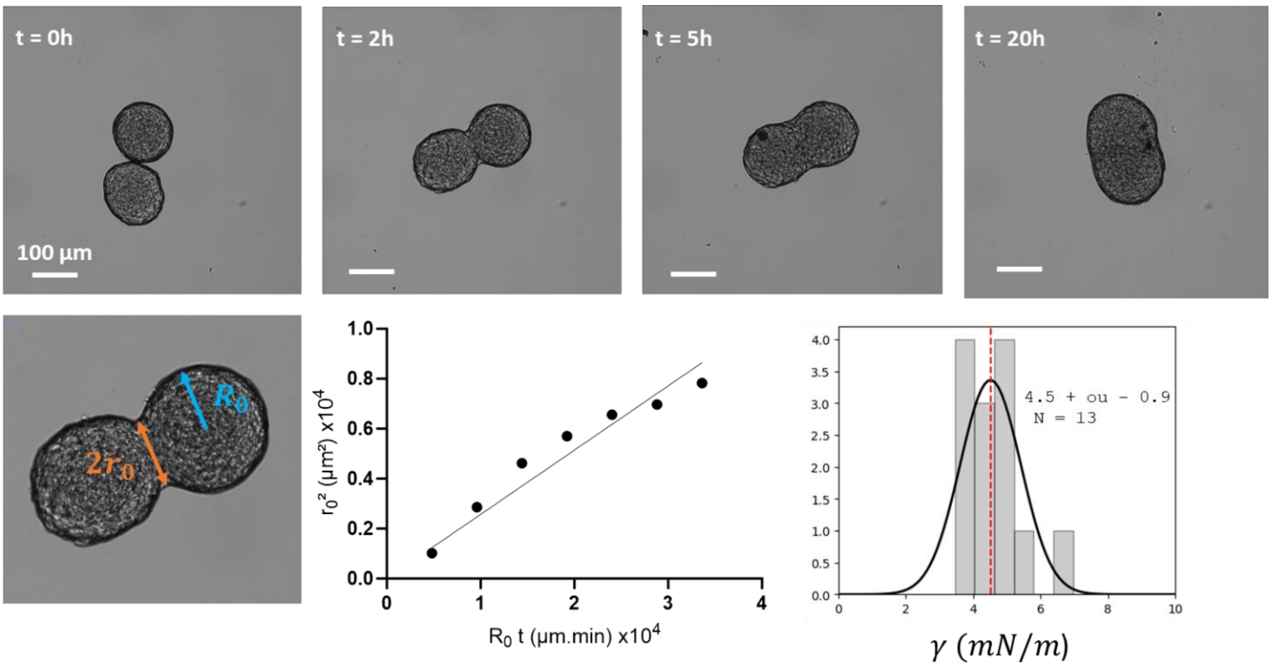


Figure SI-6- (a) Time lapse showing the fusion of two A338 spheroids placed in a culture well. (b) Temporal evolution of the size of the contact zone between the spheroids. (c) Histogram of the deduced surface tension, according to Equation (SI-1), the viscosity being fixed to $\eta = 1.2\ 10^6$ Pa.s.

As described in reference [3], the surface tension $\gamma$ was determined from the following equation, with $r_0$ the radius of the contact zone, $R_0$ the radius of the spheroid:

$$\left(\frac{r_0}{R_0}\right)^2 = 2^{\frac{2}{3}}\left(1 - e^{-\frac{t}{\tau}}\right), \qquad \text{with } \tau = 2^{\frac{2}{3}} R_0 \frac{\eta}{\gamma} \qquad \text{(Eq SI-1)}$$

The value retained for the viscosity was the one measured in the chip: $\eta = 1.2\ 10^6$ Pa.s. For $t \ll \tau_c$, $r_0^2 \approx \frac{\gamma}{\eta} R_0 t$.

### SMPs with non-circular pipette cross section permit to expose trapped micro-objects to a fluid shear stress

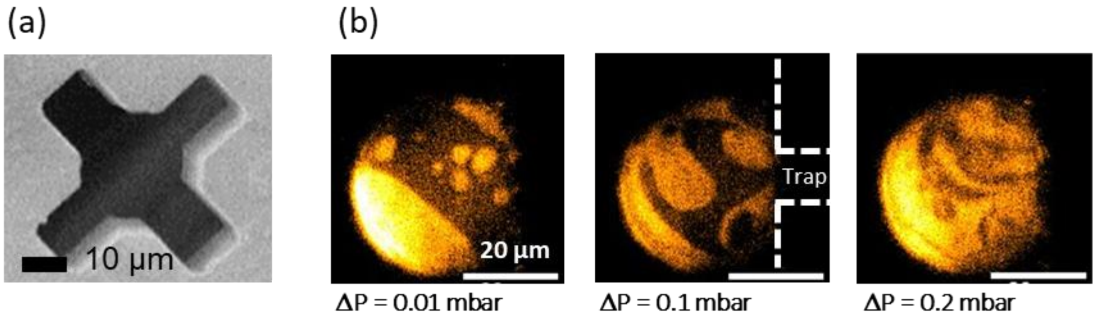


Figure SI-7- (a) SEM micrograph of a pipette with cross-shape section. (b) Giant Unilamellar Vesicle (GUV) presenting lipid domains (revealed by a selective fluorophore), blocked in a cross-shape pipette. Increasing the pressure drop applied to the pipette increased the shear stress around the GUV, and induced a merging of the domains.

Domain-forming GUVs like the one visible in Figure SI-7(b) were obtained by electroformation, with the following protocol, reproduced after reference [4]: a mixture of 32% DOPC, 40% of DSPC (1,2-distearoyl-sn-glycero-3-phosphocholine) lipids, and 28% of cholesterol . Two fluorescent dies were added to the solution: 0.1% of the fluorescent dye $DiIC_{20}$ ($\lambda_{ex}$ = 551 nm, $\lambda_{em}$ = 566nm), selective to mobile, liquid-disordered domains, and 0.1% of the fluorescent BODIPY ($\lambda_{ex}$ = 488 nm, $\lambda_{em}$ = 503nm) (selective to liquid-ordered phase) were dissolved in chloroform. A 10 µL droplet was deposited on an ITO substrate and evaporated under vacuum for 30 mn. Electroformation was realized as described in the “Preparation of Giant Unilamellar Vesicles” section, with a peak-peak Voltage equal to 2V. However, as DSPC phase transition temperature is 55°C, electroformation was done under 60°C for 2.5 hours. Slow cooling down was realized with a ramp at 5°C/min down to the room temperature.

### Control of the chemical microenvironment around trapped objects

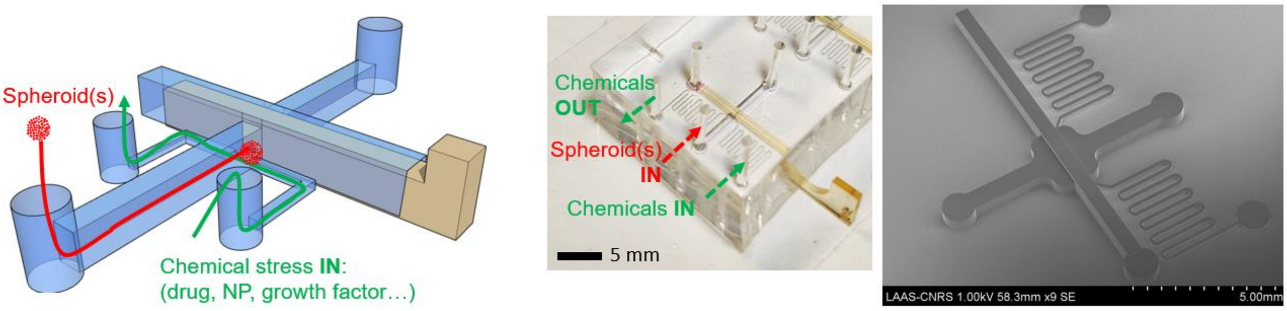


Figure SI-8- Fluidic design to capture spheroids (or vesicles) and submit them to a dynamic chemical stimulus, thanks to lateral microchannels. From left to right: schematics of the design, photo of the assembled chip (PDMS device with the SMP inserted), SEM micrograph of the design.

Supplementary Videos 4 and 5 demonstrate the operation of this chip, with fluorescent microparticles used to visualize the change of buffer surrounding the spheroids.

### Legends for Supplementary Videos

Supplementary Video 1: Aspiration of a spheroid (A338 cells) with a pressure $\Delta P = 50$ mbar and pipette radius Rp = 35µm.

Supplementary Video 2: Ejection of a spheroid (A338 cells) during retraction when the pressure is set back to zero. Scale bar 50 µm, video duration 10 min.

Supplementary Video 3: Detection of the contour and tip of the spheroid tongue during an experiment with our automated algorithm. The imposed pressure is $\Delta P = 50$mbar and the pipette radius is 35µm.

Supplementary Video 4: Real-time movie with a device for changing the environment around the spheroids. Here, 5 µm fluorescent beads are injected towards the spheroids.

Supplementary Video 5: Quick change of the medium around the spheroids in order to remove the chemicals previously added (here 5 µm fluorescent beads). Real-time movie.

Supplementary Video 6: Single Cell aspiration. Real time. Diameter of the pipettes 5µm, fluorescently labelled A338 cells, with CellTracker green fluorescent probes.

Supplementary Video 7: SMPs fabricated in glass, showing no fluorescence, which permitted imaging individual cells inside the pipette. Z-stack at constant pressure, followed by aspiration at constant Z. Glass SMPs were obtained from Femtika company. They were fabricated by selective laser etching.
